# Do goldfish like to be informed?

**DOI:** 10.1101/2024.03.17.585404

**Authors:** Victor Ajuwon, Tiago Monteiro, Mark Walton, Alex Kacelnik

**Author notes:** Correspondence should be addressed to VA, TM or AK. Both authors contributed equally to this work (Victor Ajuwon, Tiago Monteiro).

## Abstract

Most mammalian and avian species tested so far, including humans, prefer foretold over unsignalled future events, even if the information is costly and confers no direct benefit, a phenomenon that has been called paradoxical, or suboptimal choice. It is unclear whether this is an epiphenomenon of taxonomically widespread mechanisms of reinforcement learning, or if information-seeking is a dedicated cognitive trait, perhaps a precursor of human curiosity. We investigate whether a teleost fish that shares basic reinforcement learning mechanisms with birds and mammals also presents such preference, with the aim of dissociating food-reinforced learning from information-seeking. Goldfish chose between two alternatives, both yielding a 50% chance of reward 5s after being chosen. The ‘informative’ alternative caused immediate onset of either of two stimuli (S+ or S-) correlated with the trial’s forthcoming outcome (reward/no reward). Choosing the ‘non-informative’ option, instead triggered either of two uncorrelated stimuli (N1 or N2). Goldfish learned to discriminate between the different contingencies, but did not develop preference for the informative option. This shows that conditioning learning is not always sufficient, and the difference with birds and mammals supports the hypothesis that information-seeking, rather than simple conditioning, causes the paradoxical preference for unusable information shown by the latter.

## Introduction

Environmental information enables organisms to exploit opportunities, increasing the acquisition of essential resources such as food, while reducing undesirable risks. The fact that information is valuable inasmuch as it can lead to some behavioural change is part of normative views of learning (for a review see 1) and decision-making, in a range of fields including microeconomics (2), neuroeconomics (3) and behavioural ecology (4,5). However, from a mechanistic perspective, informative stimuli may become reinforcing (or appetitive) *per se*, regardless of whether this correlates with substantial gains in primary reinforcement such as food or water rewards (6–9). Notice that we use ‘reinforcing’ in the sense of exerting an effect on the probability of future expression of behaviour, while ‘appetitive’ has a stronger connotation of goal-seeking and/or being hedonically influential. Bridging the gap between the normative rationale and empirical results from studies examining the reinforcing effect and hedonic value of informative stimuli has widespread interest for behavioural biology and comparative cognition.

Some experimental observations and theoretical models support the view that individuals may ascribe value to stimuli because they are informative, independent of associated instrumental utility (10–13). In particular, in an experimental protocol often referred to as ‘paradoxical choice’, or ‘non-instrumental information-seeking’, humans (14–18) and subjects from other mammalian (19–24), and avian (25–29) species, seek information about forthcoming outcomes (i.e. food/no food), even though in the experimental protocols the information cannot be used to modify outcomes, and may come at considerable cost (see also “observing responses” 8). In one example, starlings (*Sturnus vulgaris*) significantly preferred an option that gave food once every 20 trials over an alternative that gave food once every 2 trials, when the only additional difference was that in the leaner option the outcome was signalled during the 10 seconds of waiting between choice and outcome (Vasconcelos et al, 2015). In such protocols, subjects are typically presented with two options which deliver food or liquid rewards probabilistically, a fixed delay after the choosing response. In the informative option (Info) ether of two discriminatory stimuli are immediately presented after each choice, one paired with forthcoming reward and the other with no reward. In the uninformative option (NoInfo), outcomes are not differentially signalled, and food is either delivered or not once the delay has lapsed. The probabilities of food/no food vary depending on the implementation. Critically, subjects cannot exploit the information in the Info alternative because it is supplied during a waiting delay after they have chosen.

Some authors argue that the observed preference for the informative option reflects intrinsically motivated information-seeking mechanisms concerning future events of biological significance (7,11,14,15). Neuroscientific studies on macaques (e.g., 30,31) have helped to advance the notion that preferences in such tasks reflect non-instrumental information-seeking that is an analogue (or homologue) to human curiosity – an intrinsic drive to investigate and learn – reflecting psychological mechanisms aimed at resolving uncertainty (14,17,32) or filling ‘information gaps’ (33). This view is supported by evidence in both humans (15,34,35) and monkeys (20,21,35) showing that the same midbrain dopamine neurons can encode both information gain and primary rewards such as food. However, there are alternative interpretations.

Studies on pigeons have helped to characterise a range of experimental factors that influence preference in paradoxical choice tasks (8,26,36,37), and have advanced an alternative mechanistic account of observed preferences that do not posit a sensitivity to uncertainty reduction *per se*. According to this account, Pavlovian stimuli paired with positive outcomes become appetitive, conditioned secondary reinforcers, while stimuli paired with negative outcomes (such as omission of food delivery) are weakly inhibitory or ignored all together (8,28,36,38). If signals for positive outcomes cause a greater increase in probability of the preceding choice than the decrease caused by signals for negative outcomes, then the net result can differentially favour the ‘informative’ option, just by the asymmetric effect of secondary reinforcement. This is an appealing, parsimonious interpretation that does not rely on uncertainty-related cognition. Similarly, a study on humans has suggested that ‘good news’ stimuli derive value by allowing subjects to hedonically savour forthcoming outcomes while they wait for them (16). Again, this requires that the ‘savouring’ effect in the presence of signals for positive outcomes is stronger than the aversive ‘dreading’ effect of a foretold failure to receive reward.

From a functional perspective, some authors argue that the psychological mechanisms responsible for paradoxical choice may reflect unavoidable cognitive constraints that are also implicated in pathological human gambling (16,39–42). This hypothesis is not entirely appealing, because it neglects the question of why such maladaptive behaviour would not be eliminated and/or modified by natural selection.

In contrast, other authors aim at functionally explaining the observations by relating them to hypothetical scenarios in the environment of evolutionary adaptation (i.e. not in the lab). A foraging analysis by Vasconcelos (29) et al. (see also 25,43) argues that preferences for advanced information leading to submaximal outcomes in the lab could maximise reward rates in natural situations, because in the wild information is most often useable, unlike in the experimental protocols. Thus, adaptive mechanisms evolved to surmount common foraging challenges result in paradoxical observations in the laboratory (44). The relatively recent discovery of robust preference for reward-predictive stimuli in paradoxical choice with rats (19,22,23) supports this notion: the prevalence of the phenomenon in distantly related taxa is consistent with the view that that information-seeking is an adaptation to common, widespread ecological challenges.

Though theoretical advances are being made in functional and mechanistic understanding of apparent information-seeking behaviour, so far studies have been restricted to a relatively small set of mammal and bird species, primarily macaques, starlings and pigeons. In contrast, reinforcement processes are known to be shared by a wider range of taxa. If it could be shown that the paradoxical observations are replicable in all species endowed with ordinary reinforcement learning – that is those whose behaviour is driven by conventional reinforcers rather than putative abstract variables like uncertainty – then the need to invoke a direct appetitive influence of information per se would be weakened.

We address these issues here by implementing the paradoxical choice protocol in goldfish (*Carassius auratus)* – a species that is phylogenetically distant to those that have so far been tested. Goldfish are a suitable starting point to explore the widespread prevalence of paradoxical choice because, though relatively under-utilised, they are an established model in cognitive research (Bitterman, 1975; Salena et al., 2021), including in associative learning tasks (45–47), and their behaviour can be reinforced by conditioned stimuli (48), making them a good candidate to examine the subjective value of informative stimuli. Goldfish also diverge dramatically in brain architecture from the mammalian and avian species so far tested (49), and thereby provide an opportunity to test the conjecture that derived neocortex - like structures are critical to information-seeking behaviour (31).

## Methods

### Subjects

Eight goldfish ranging in size between 7 - 10cm, (age and sex unknown) participated. They were obtained from a local commercial supplier (Goldfish Bowl, Oxford, UK) and housed in groups of two or three, in holding aquaria (60 x 35 x 31 cm; *length* x *width* x *height*) where they had access to a rock shelter, pebbles and artificial plants. Individuals participated in experiments 5 times a week on weekdays and were fed *Fancy Goldfish Sinking Pellets* during sessions. This diet was supplemented with spinach following experiments on the last day of the working week and bloodworms the day after (subjects were not directly fed on Sundays). Fish were kept under a 12:12 h light: dark cycle using fluorescent lights. Water was maintained at a minimum of 21℃ using an internal heater and independent thermometer (pH: 8.2; ammonia: 0 ppm; nitrite: 0 ppm; nitrate: max. 30 ppm). Partial water changes were conducted at the end of each week and internal filters were cleaned every month. Each holding tank was aerated using an air pump.

For each daily session, fish were transported to their experimental tank and then back to their holding tank in a plastic jug. At the start of each day ∼20L of water from the holding tanks were transferred to the experimental tanks in order to keep the environmental conditions as constant as possible. The experimental tanks were cleaned at the end of each week. The animals had different levels of previous experience in unrelated protocols.

### Apparatus and task control

Experiments were run using *GoFish*, an open-source experimental platform for closed-loop behavioural experiments on aquatic species (46). The use of *GoFish* brings a similar level of automation and control to that found in previous paradoxical choice experiments with other species, aiding interspecies comparisons and reducing potential experimenter-induced bias.

The experimental set-up constituted a rectangular tank (60 × 30 × 36 cm; *length* x *width* x *height*) with 15 cm depth of water; a 17’’ LCD computer screen (1920 × 1080; 60 Hz) for stimulus presentation; two custom-made pellet dispensers placed either side of an opaque acrylic divider in a y-maze configuration; an overhanging USB camera (1280×720 resolution); and an aquarium light (**Figure 1a**). Each reward delivery consisted of one *Fancy Goldfish Sinking Pellet.* Experiments were run in parallel in two experimental tanks with 4 fish randomly assigned to each at the start of the experiment.

**Figure 1.**
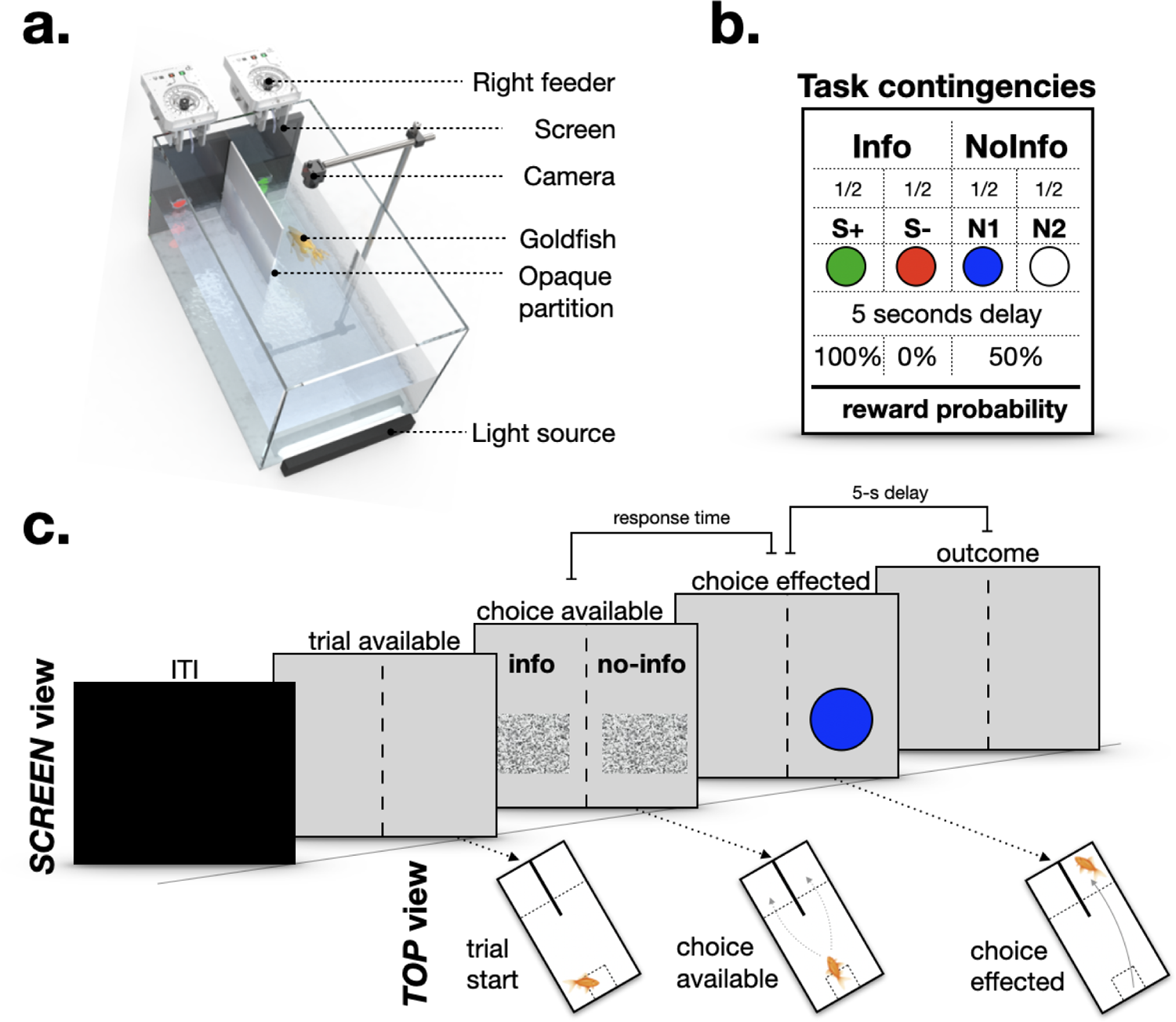
Exploring preference for advanced information in goldfish. **a**. The *GoFish* operant chamber includes a screen for stimulus presentation, two pellet dispensers placed either side of an opaque acrylic divider in a y-maze configuration, an overhanging camera and aquarium light. **b**. In Phase 1 of the experiment each subject chose between two options: in Info the forthcoming reward outcome was signalled immediately post-choice, in NoInfo stimuli were not outcome-predictive. **c**. Illustrative example of a Phase 1 trial showing a subject initiating the trial and choosing the non-informative option.

Task control was automated and implemented using a custom *Bonsai* workflow (50). Task contingencies (i.e., stimulus presentation and reward delivery as a function of behaviour) were dependent on fish being within specified regions-of-interest (ROIs) in the experimental tank, and were controlled using real-time video detection of fish movement. Within each tank three ROIs were defined: a ‘start zone’, ‘left choice zone’ and ‘right choice zone’. These zones were not defined by physical boundary markings but were delineated on the video feed corresponding to fixed areas within the experimental tank. To detect fish presence in the ROIs, the following method was used. Video of the experimental tank was recorded at approximately 33 fps and converted to HSV colour space. A HSV range was applied to detect the fish. The pixels of the resulting binarized frames from each ROI were summed continuously. Fish entries into the zones were recorded when the summed ROI pixel value exceeded a pre-set threshold (ROI thresholds were adjusted to each subject prior to the onset of the experiment). Thus, subjects were able to control the progress of trials and make choices via their position in the tank. We also used colour thresholding to continuously track the centroid of each fish throughout experimental sessions (for further details see 46).

### Pre-training

Pre-training consisted of three phases: (i) experimental tank acclimatisation, (ii) choice zone training, for subjects to learn that swimming into either the left or right choice zone (outside of the inter-trial interval, ITI) could be followed by food delivery, and (iii) start position training, for subjects to learn that a start position had to be entered before entry into choice zones was reinforced. Advancing through the phases depended on the individual subject’s performance. For further details see ‘Pre-training’ in (46).

### Phase 1

We used a trial-based chain procedure. Every fish was presented with 2 daily sessions of 24 trials for 45 days. There were 2 kinds of trials: 2-option choice trials and 1-option forced trials. The proportion of each of these trials within a session changed as the experiment progressed (**Supplementary Table 1**) but for the first 15 days subjects were exposed only to forced trials. Subsequently, where a session was composed of both types of trial, they were randomly intermixed. Trials were separated by an ITI (drawn from a uniform distribution: min = 20 s; max = 40 s) during which the screen was black and behaviour had no consequences. The end of the ITI was signalled by a grey screen and from this moment on, entering the start position would trigger the presentation of a visual white noise stimulus in the half screen facing one of the two choice zones (1-option forced trial) or both choice zones (2-option choice trial). Swimming into either choice zone with the white noise stimulus present led to the substitution of the white noise stimulus by a geometric shape visual stimulus (S+, S-, N1 or N2, see ‘Stimuli’ section below) which remained present for 5s, until the end of the trial.

### Paradoxical Choice Experiment

There were two options (Figure 1b), which could either be presented alone (1-option forced trial) or simultaneously (2-options choice trial). Swimming into the choice zone of the informative option (Info) resulted with equal probability in either of the sequences “S+ ⇒ wait for 5s ⇒ Food”, or “S- ⇒ wait for 5s ⇒ Nothing”, thereby reliably informing the subject of the forthcoming outcome. Swimming into the non-informative (NoInfo) choice zone, on the other hand, resulted with equal probability in either of two sequences: “N1 ⇒ wait for 5s ⇒ 50% either Food or Nothing” or “N2 ⇒ wait for 5s ⇒ 50% either Food or Nothing”; therefore, in NoInfo neither cue informed the subject of forthcoming outcome, but reward probability was the same as when choosing Info (an example choice trial is shown in Figure 1c). The sides of the Info/NoInfo options were consistent for each individual but counterbalanced across subjects.

### Phase 2

After 30 days of being presented with choices between Info and NoInfo, subjects were presented with randomly intermingled choices between the following pairs of stimuli: S+ vs either N1 or N2, and S- vs either N1 or N2. The aim of this phase was to test whether the fish had learned the contingencies of the terminal links in the chain trials. There were 2 daily sessions of 24 trials per day, which lasted for 5 days. Trials were preceded by an inter-trial-interval (ITI; drawn from a uniform distribution: min = 20 s; max = 40 s) where the screen was black and behaviour had no consequences. The end of the ITI was signalled by a grey screen and from this moment on, entering the start position triggered the presentation of S+ or S- in one choice zone and N1 or N2 in the other. Swimming into a choice zone led to the reward outcome for the chosen stimulus according to the same contingencies as in Phase 1, followed by the onset of the ITI.

### Stimuli

In Phase 1 of the main experiment we used a monochromatic white noise rectangle (13.5 x 12 cm, Gaussian: mean=0, variance=10) on either the right or left half of the screen (or both) to indicate whether the choice zones were primed to advance the protocol upon entering. In the pre-training phase, the white noise stimulus was used to signal that reward delivery was contingent on fish swimming to a specific location in the tank.

The outcome-predictive cues were 4 coloured circles presented on the screen, each paired with a reward probability (1, 0, or 0.5, for S+, S- and N1/N2 respectively, with colours counterbalanced across subjects). These were red, green, blue, and white circles, 3.5 cm in diameter on a grey background, with centres positioned 5 cm from the bottom of the tank and 7 cm from each side wall. We chose colours that have been physiologically (51) and behaviourally (46,52) proven to be discernible by goldfish.

### Data analysis

To examine whether subjects learned the contingencies of each outcome- predictive stimulus we quantified fish spatial occupancy in a 4s time window preceding reward outcomes (excluding 0.5s after choice and 0.5s pre outcome) using entropy (a measure of randomness) as a ‘movement metric’ to parameterise the spatial distribution of subjects from an input matrix capturing this 4s window. Entropy (and therefore our movement metric) is given by

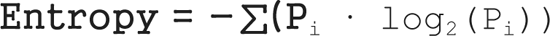

as implemented in the function `Entropy` (53), where *P* are matrices containing the normalised histogram (3 cm^2^ bins) counts for each stimulus type, S+, S-, N1 and N2, with N1 and N2 pooled together, since they have identical reward contingencies. Data processing and analysis was carried out in MATLAB 2022b. Statistical analyses were conducted in RStudio (v1.2.5033; 54). Analyses were carried out on data from the last 30 days of *Phase 1* of the experiment, in which forced and choice trials were intermixed, and all 5 days of *Phase 2*. Response time and occupancy data are from the last 5 days of Phase 1. Due to accidental misalignment of the camera, 13 out of the 80 sessions were omitted from analyses of occupancy data as detailed in **Supplementary Table 2**. Visual inspection of data trends indicate that this loss did not have the potential for substantially change any of the conclusions. To normalise the residuals in statistical analyses, response time data were log transformed, while choice proportion data were arcsin square-root transformed. A type-1 error rate of 0.05 was adopted for all statistical comparisons. A Bonferroni correction was applied for paired pairwise comparisons.

### Ethics statement

All experiments were conducted at the John Krebs Wytham Field Station of the University of Oxford, and approved by the Department of Biology Ethical Committee, University of Oxford (Ref. No. APA/1/5/ZOO/NASPA/Ajuwon/Goldfish). They were carried out in accordance with the current laws of the United Kingdom. Animals were cared for in accordance with the University of Oxford’s “gold standard” animal care guidelines. All experimental methods were non-invasive, and no food restriction was necessary.Maintenance and experimental protocols adhered to the Guidelines for the Use of Animals in Research from the Association for the Study of Animal Behaviour/Animal Behavior Society (55). On completion, the fish were reintroduced into holding tanks and retained for further experiments.

## Results

### Phase 1

#### No evidence goldfish prefer advanced information in paradoxical choice

Our subjects showed no evidence of preference for the option in which outcomes were signalled between choice and trial end (Figure 2a). Over 30 days individual subjects were presented with over 500 choices: A one-way repeated measures ANOVA with day as a within-subject factor and proportion of choices for Info as the response variable showed no significant effect of day (F_29,203_ = 1.00, *P* = 0.471), indicating that on average the subjects did not develop a preference for either option.

**Figure 2.**
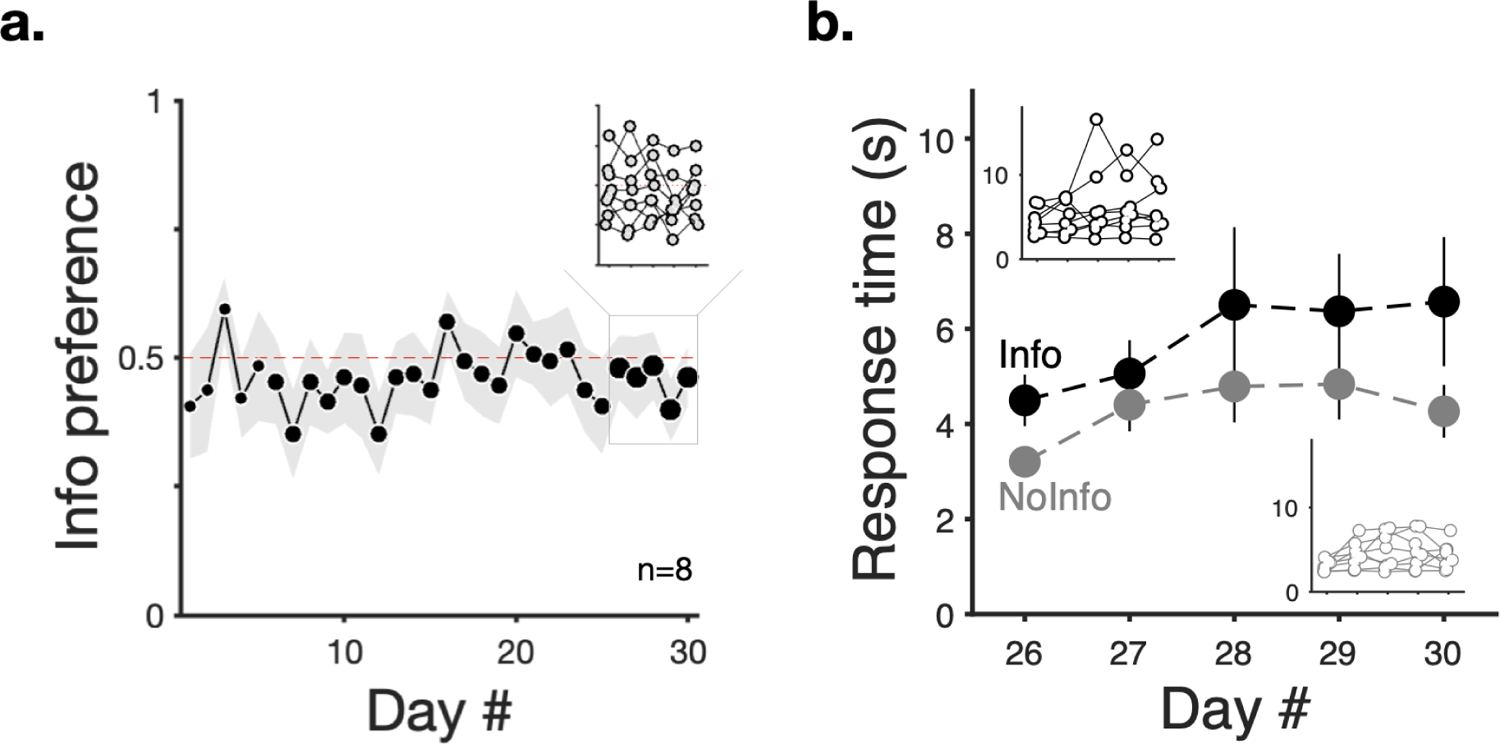
Goldfish preferences and response times for the informative alternative. **a.** Mean proportion of choices for the informative option in 2-option choice trials (n=8; grey patch depicts ± SEM). Marker size is proportional to the number of 2-option choice trials - for details see *Supplementary Table 1*. **b.** Mean response times (n=8; in seconds ± SEM) for responding to Info (black) and NoInfo (grey) when encountered in single-option trials during the last 5 days of *Phase 1*. In both panels insets show individual animal means for the last 5 days of the experiment.

In addition to 2-option choices, during this period subjects were also presented with 1-option forced trials. In these trials we recorded the time latency from subjects starting a trial to proceeding to the single option presented to them; the ‘response time’. Response times often are a more sensitive metric of valuation than choice proportion in simultaneous trials, with shorter response times in one-option encounters associated with greater preference in choices (56–58). We examined response times in forced trials with either alternative during the last 5 days of the experiment. Figure 2b shows that on average individuals took longer to respond in trials with the informative option than in trials with the non-informative option. This was reflected in a two-way repeated measures ANOVA with day and option as within-subject factors and response time as the response variable which showed a significant effect of day (F_4,28_ = 4.95, *P* < 0.01), and option, (F_1,35_ = 9.09, *P* < 0.01), but no interaction (F_4,35_ = 0.217, *P* = 0.927). Consistent with previous studies (e.g., 57,59) latencies in forced trials and proportion of choices were related. **Supplementary Figure 1** shows a negative correlation between individuals’ strength of preference for the informative option and the difference in response time between both options (individuals with shorter reaction times for a given option in forced trials show greater preference for that option in choices).

#### Post-choice goldfish occupancy shows discrimination of stimuli signalling future outcomes

To investigate whether the lack of preference between the options showed in choice trials was due to a putative inability of goldfish to learn the contingencies associated with either option, we examined subjects’ behaviour towards the outcome-predictive stimuli in each alternative. We focused on goldfish locomotion, for the last 5 days, during a 4s portion (cropping off 0.5 s after choice and 0.5 s pre-outcome) of the delay period preceding reward outcomes. This shows that goldfish discriminated between cues: proximity to the sites of reward delivery varied according to each cue’s reward contingency (recall that reward probabilities were S+ 100%; N1/N 50%; S- 0%). When reward was forthcoming (Figure 3a**,b** left panels) or potentially forthcoming (Figure 3a**,b** centre panels) subjects generally remained in the choice zone, close to the pellet dispenser. When reward was not forthcoming (Figure 3a**,b** right panels) subjects actively swam away from the choice zone, and pellet dispenser, a trend shown for each subject (**Supplementary Figure 2**).

**Figure 3.**
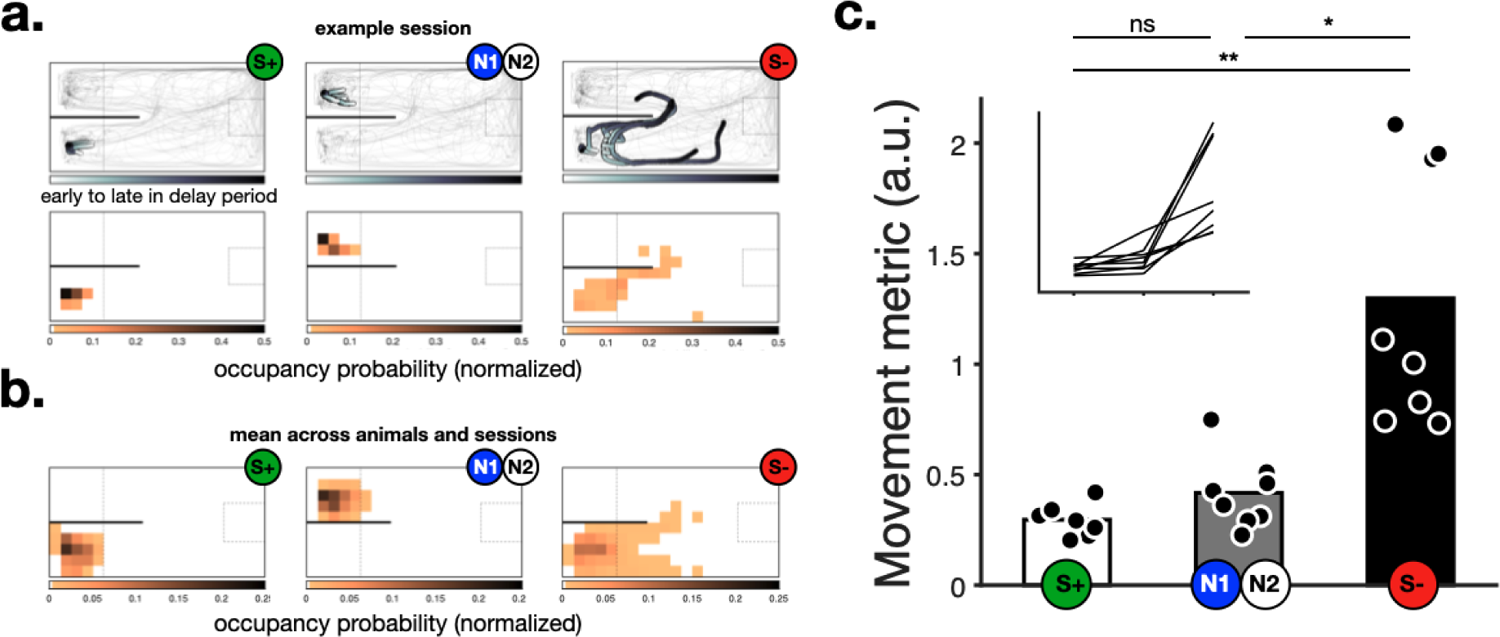
Goldfish spatial location varied according to cues’ reward contingencies. **a. Top row**: trajectories of a representative animal and session during the post-choice delay period, split by trials in which the animal was presented with S+ (100% reward, left), N1 or N2 (50% reward, middle), or S- (0 % reward, right). Darker trajectory shading shows the subject’s location further into the delay period. Thin grey lines show the trajectories for the entire session for reference. **Middle row**: Corresponding normalised density heat maps of fish occupancy, darker shading indicates greater occupancy. **b.** Same as above but averaged across all animals and days (n=8 animals, 5 days each). For illustrative purposes, responses to Info stimuli appear on the left and NoInfo stimuli on the right, though this was counterbalanced across subjects **c**. Movement metric computed as the entropy of subjects’ spatial distributions (see *Methods* for details), where larger values correspond to more movement. Markers show individual animal means and bars the group averages. Inset shows individual trends connecting individual data points shown in the main figure (same scale). *p<.05, **p<.01.

To quantify the behavioural responses of subjects to post-choice reward-predictive stimuli, we calculated the entropy of subjects’ spatial probability distributions during the same post-choice delay (i.e. cropping off 0.5s after choice and 0.5s pre outcome). Higher entropy values indicate subjects wandered over a larger portion of the experimental tank during this period. Data pooled from the last 5 days show that fish spatial occupancy during cue presentation corresponded to the probability of food delivery (Figure 3c), indicating cue discrimination. The entropy of spatial distributions (mean ± s.e.m.) was lowest when S+ was presented (0.30 ± 0.024), higher when N1/N2 were presented (0.42 ± 0.058) and greatest when S- was presented (1.30 ± 0.21), reflecting that subjects spent more time near the site of food delivery when reward probability was greater. A one-way repeated-measures ANOVA on these data with stimulus as a within-subject factor and entropy as the response variable revealed a significant effect of stimulus (S+, N1/N2, S-; F_2,14_ = 18.48, *P* < 0.001). Post-hoc pairwise comparisons showed that entropy of spatial distributions was significantly different between S+ and S- (*P* < 0.01), S- and N1/N2 (*P* < 0.05), but not between S+ and N1/N2 (*P* = 0.165), though subjects did show greater adherence to the site of reward delivery when S+ was present compared to N1 or N2. Overall, these results indicate that subjects did discriminate the contingencies, and potentially developed an aversive response to S-. Interestingly, previous work in pigeons has emphasised the appetitive effect of S+ over the aversive effect of S- (36,37).

### Phase 2

#### Goldfish choices between reward-predictive stimuli suggest biased weighting of reward probability

The results from Phase 1 show that subjects were indifferent between a site where their choice led to either S+ or S- and a site where the post choice stimuli (N1 and N2) did not predict the trials’ outcome, but their post-choice behaviour showed discrimination between the stimuli themselves. This is evidence for learning the contingencies, but does not prove that goldfish can express such learning in their choice of feeding site. To corroborate whether they could express stimulus valuation as site preferences, in Phase 2 trials started directly with presentation of pairs of stimuli that in Phase 1 had been experienced after their choices (**Supplementary Figure 3a**).

When choosing between S+ and either N1 or N2, subjects were, on average, indifferent, while when choosing between S- and either N1 or N2, they had a clear preference for the latter. A two-way repeated measures ANOVA, with stimulus and day as within-subject factors and proportion of choices for S+/S- as the response variable revealed a significant effect of stimulus (S+ or S-; F_1,35_ = 85.19, *P* < 0.0001), but no significant effect of day (F_4,28_ = 0.87, *P* = 0.0494), and no significant interaction (F_4,35_ = 1.13, *P* = 0.357; **Supplementary** Figure 3b). Specifically, subjects showed no preference for S+ against N1/N2 (S+ choices: 46%, ± 0.072 s.e.m; t7 = −0.63, P = 0.551), but preferred N1/N2 against S- (S- choices: 17%, ± 0.029; t7 = −8.97, P < 0.0001). Taken together, these results indicate that subjects could express their knowledge about S- by choosing away from it. This is consistent with results from Phase 1, where there were similar locomotory responses to S+ and N1/N2, compared to S-, from which subjects moved away.

## Discussion

In contrast to the most common vertebrate species previously tested (humans, monkeys, rats, pigeons and starlings), here goldfish did not display a preference for signals foretelling probabilistic reward outcomes in the paradoxical choice task (Figure 4), though they did discriminate between their contingencies. Our results have implications for our understanding of the psychological mechanisms underlying information-seeking behaviour, as well as its functional significance and evolutionary origins.

**Figure 4.**
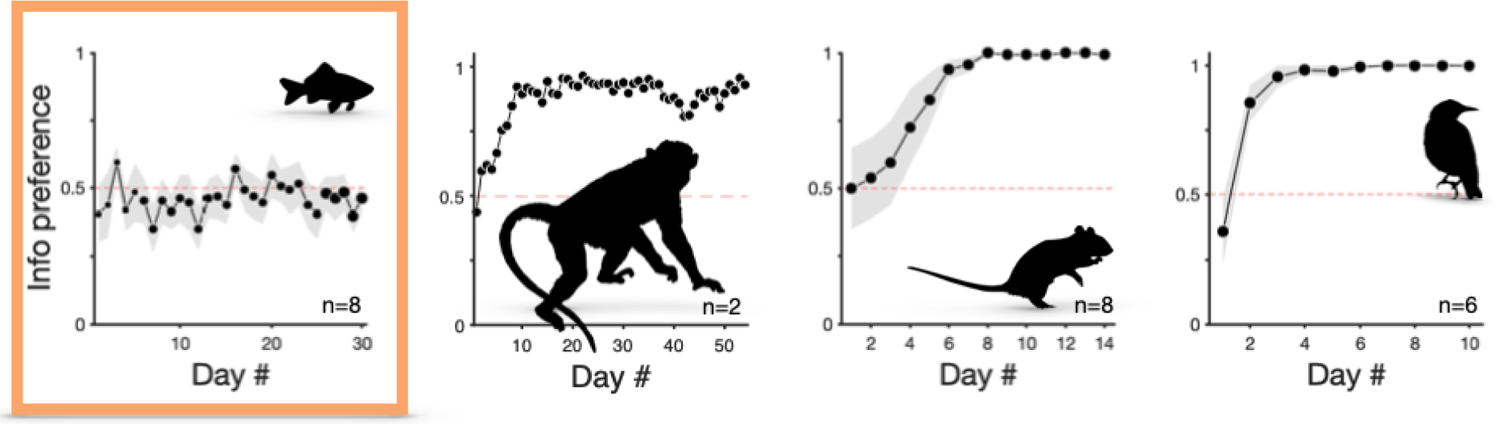
Preference for the informative alternative across 4 different species in similar paradoxical choice protocols. From left to right: Goldfish (*Carassius auratus*; n = 8) from the present study (same as Figure 2a). Rhesus macaque (*Macaca mulatta;* n = 2). Adapted with permission from (21) - Figure 1b. Rats (*Rattus norvegicus*, n = 8). Adapted with permission from (19) - Figure 3. European starlings (*Sturnus vulgaris*, n = 6). Adapted with permission from (29) - Figure 2. Markers show group means and shading the s.e.m. The orange box highlights that the data are from the present study. Animal silhouettes from https://www.phylopic.org/ under Creative Commons licences.

Key to the interpretation of our results is the ability of subjects to discriminate between the outcome-predictive cues that distinguish the informative and uninformative alternatives. Two independent pieces of evidence suggest that the goldfish in our study did discriminate between the outcome predictive stimuli. First, during the delay period between choice and outcome, when cues were present in the main experiment, subjects’ proximity to the site of reward delivery showed that they treated the cues according to their reward contingencies: the greater the likelihood of reward, the more time spent close to the site of food delivery (Figure 3).

Similar discrimination has been shown in rats, but the latter did show a preference for the option offering advanced information (19,23). Second, in Phase 2, we presented choices between signals that predicted the trial’s outcome (S+ or S-) and signals that did not (N1 or N2). In Phase 1 these cues had only been presented after the choice of option, while in Phase 2 fish could choose directly between them. The purpose of these tests was to establish whether choice indifference in the main experiment could be due to lack of post-choice signals discrimination or an inability to express valuation as choice, as opposed to lack of preference. In this phase, the fish were indifferent between signals for sure or 50% chance of reward, but preferred signals for 50% reward over a signal for reward absence. Interestingly, this result differs partly from results in starlings, that showed both a preference for sure over 50% chance reward and a preference for 50% over sure omission (Vasconcelos et al, 2015). Indifference between 100% and 50% reward signals was unexpected, but could be due to a ceiling effect: if valuation is a strongly concave function of reward probability, the differential attractiveness of medium and high probability contingencies may be undetectable. Our result is reminiscent of the observation reported by Stagner and Zentall (2010), who found that pigeons preferred partial over continuous reinforcement and interpreted it as a suboptimal extreme form of risk propensity. This remains an interesting research topic in its own right.

Our protocol retained as much as possible the critical features of the task used in other species. Furthermore, unlike studies where subjects incur reward losses by preferring the informative option (24,25,29,60–63), to avoid the possibility of masking preference for information due to sensitivity to loss, in our study the informative and uninformative options were equally profitable.

From a procedural perspective, task parameters other than profitability (i.e. Reward/Delay; 64) can influence the strength of preferences in paradoxical choice tasks (for reviews see 36,37). These include the salience and modality of outcome predictive stimuli (19,29,40,65), the length of delay between choice and reward outcomes (longer delays lead to greater *Info* preference: (16,22,60,66), the proportion of forced trials (more forced trials increase preference: (27) and the action subjects must make to express preference (67,68). Therefore, it is possible that our results depend on specific experimental parameters, with a particular issue being that the length of delay between choice and reward outcomes may have been too short to elicit a preference for the informative option. However, *a priori* it is not obvious why such details would have masked an underlying preference for information, for several reasons. In the first place, similar parameter sets have generated robust preferences in other species. We designed our experiment so as to minimise the potential influence of specific parameter values, by using counterbalanced stimuli to which goldfish are known to respond (46) and selecting a delay duration (5s) long enough to elicit preferences in monkeys (20,21,i.e., >2.5s; 30,35) but shorter than the 10s used in rat, starling and pigeon studies, to reduce the possibility of task disengagement in the freely swimming fish. Furthermore, our subjects underwent training with 1-option forced trials before 2-option choice trials were introduced, and forced trials were interspersed throughout the main experiment, reducing the possible influence of side biases. These considerations, taken together with the fact that subjects learnt to discriminate the cues associated with each option under our set of experimental parameters, lead us to believe that our results probably do reflect genuine species differences between goldfish and other vertebrate models in sensitivity to advanced information about forthcoming outcomes.

Why (mechanistically and functionally) may such species differences exist? One possibility concerns the effects of conditioned reinforcement. One mechanistic explanation for learned preferences for advanced information argues that subjects select the informative option because ‘good news’ signals anticipating forthcoming rewards acquire reinforcing properties by association with outcomes of positive valence (8,19,28,69). If any putative inhibitory effects of signals for ‘bad news’ in the informative option are relatively small or ignored by subjects, overall, subjects would form stronger associations between choice and reward in the informative option, but these effects may be quantitatively different between species. Rat preferences for the informative option have been shown to be less robust than other species’ (22,23,70), and it has been suggested that this is because rats may be more sensitive to conditioned inhibition through signals for bad news in the informative option than birds (70–72). It is therefore possible that goldfish do not prefer the informative option because the signal for bad news acquires inhibitory properties powerful enough to eliminate any reinforcing effects of the signal for good news. Some of our results indicate that this may be the case. In Phase 1, when presented with S-, subjects swam away from the stimulus (Figure 3;(see also 43)), and in Phase 2 subjects rarely selected S- when it was presented alongside other stimuli (**Supplementary** Figure 3), suggesting that stimuli for reward omission can acquire aversive properties which could inhibit Info preference. Manipulations involving the removal of S- (e.g., 19,73,74) could help to elucidate the extent to which conditioned inhibition plays a role in goldfish preferences for advanced information. Subjects in our study learnt the task contingencies, showing appropriate responses to outcome-predictive stimuli, but they did not develop a preference for informative stimuli foretelling good and bad news equally often. This suggests that explanations of paradoxical choice based exclusively on secondary conditioning, that is, excluding any role for information seeking per se, may not be sufficient to account for preferences observed in other species.

Another candidate explanation for lack of preference for informative stimuli in goldfish is that they lack sensitivity to reductions in uncertainty which is present in mammals and birds. If confirmed, this would trigger a different paradox: increased cognitive or neural sophistication, allowing handling of uncertainty, could cause loss of foraging performance; species sophisticated enough to find information rewarding *per se*, may choose suboptimally when allowed to seek for unusable information. All of this is, of course, speculative at this stage.

Functionally, the lack of preference for advanced information in goldfish may reflect a difference in foraging mode compared to other species so far tested. Vasconcelos et al. (29) argued that preferences for a lower pay-off alternative providing advanced non-instrumental information in the lab may reflect adaptive mechanisms that are rate maximising in natural circumstances, where information about forthcoming rewards will likely be usable. They envisaged that decision-makers could use such information to avoid unnecessary opportunity costs incurred while pursuing prey. The relevance of this line of reasoning to particular species, including carp that are primarily suction feeders and scavengers, can only be established through field research in ecological circumstances, but it is enough to sustain the view that paradoxical behaviour in the lab does not justify the epithet of ‘suboptimal’. Further, goldfish (like experimental rats and mice, but not starlings), are laboratory bred and reared, and this may have modified their behaviour with respect to wild ancestors growing in ecological circumstances. We acknowledge that they may have low representativeness respect to the whole fish community. If, however, further research proves that secondary conditioning is present across multiple fish species that do not show preference for advanced informative signals, while a variety of birds and mammals continue to confirm the current picture, questions about phylogenetic emergence would acquire relevance and importance: are there substantial and consistent differences in post-reptilian lifestyles that make information-seeking qualitatively more influential than it was before the split of tetrapods from the ancestral vertebrates trunk? If so, what are these differences? If nothing else, these ‘known unknowns’ add weight to the view that cognitive research benefits from escaping the use of a narrow set of laboratory species to potentiate a comparative, adaptive perspective.

## Acknowledgements

We thank Adelaide Sibeaux for technical advice and husbandry training, Theresa Burt de Perera, Cait Newport, Christine Soper and all the Wytham Field Station staff for logistical support, and Bruno Cruz for assistance with the Bonsai workflow and the Champalimaud Research Hardware and Software platform for building our automatic feeders. Many conversations with our colleagues and friends, especially Marco Vasconcelos and Tom Zentall, have left a strong imprint on our views on the topic of this paper.

## Author contributions

VA, TM MW and AK, conceptualised and designed the experiment; VA and TM collected the data; VA, TM and AK analysed the data; VA and TM wrote the first draft of the paper; all authors edited and reviewed the manuscript.

## Data and Code availability

Data and code for analysis are available upon request.

## Supplementary Materials

**Supplementary Figure 1.**
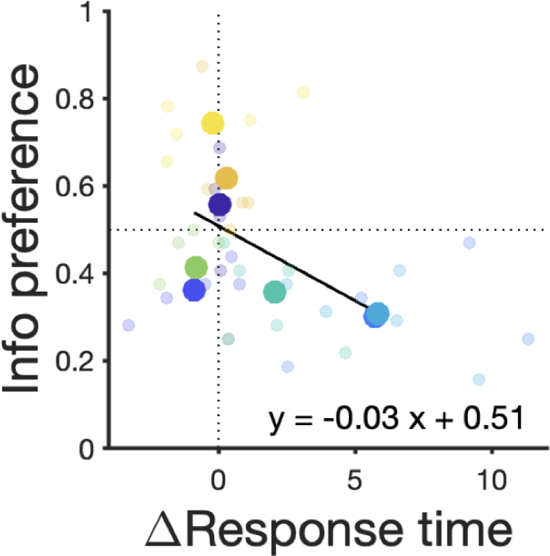
Preference for the informative option (Info) is correlated with response times. Mean proportion (n=8) of choices for the last 5 days of *Phase 1* of the Experiment 2-option choice trials as a function of the mean difference (delta) between Info and noInfo response times in 1-option forced trials (large markers). Smaller makers depict individual means across sessions. Solid black line shows a Deming regression (Estimated model: y = −0.03x + 0.51; slope 95% CI = [−0.07 to 0.33]).

**Supplementary Figure 2.**
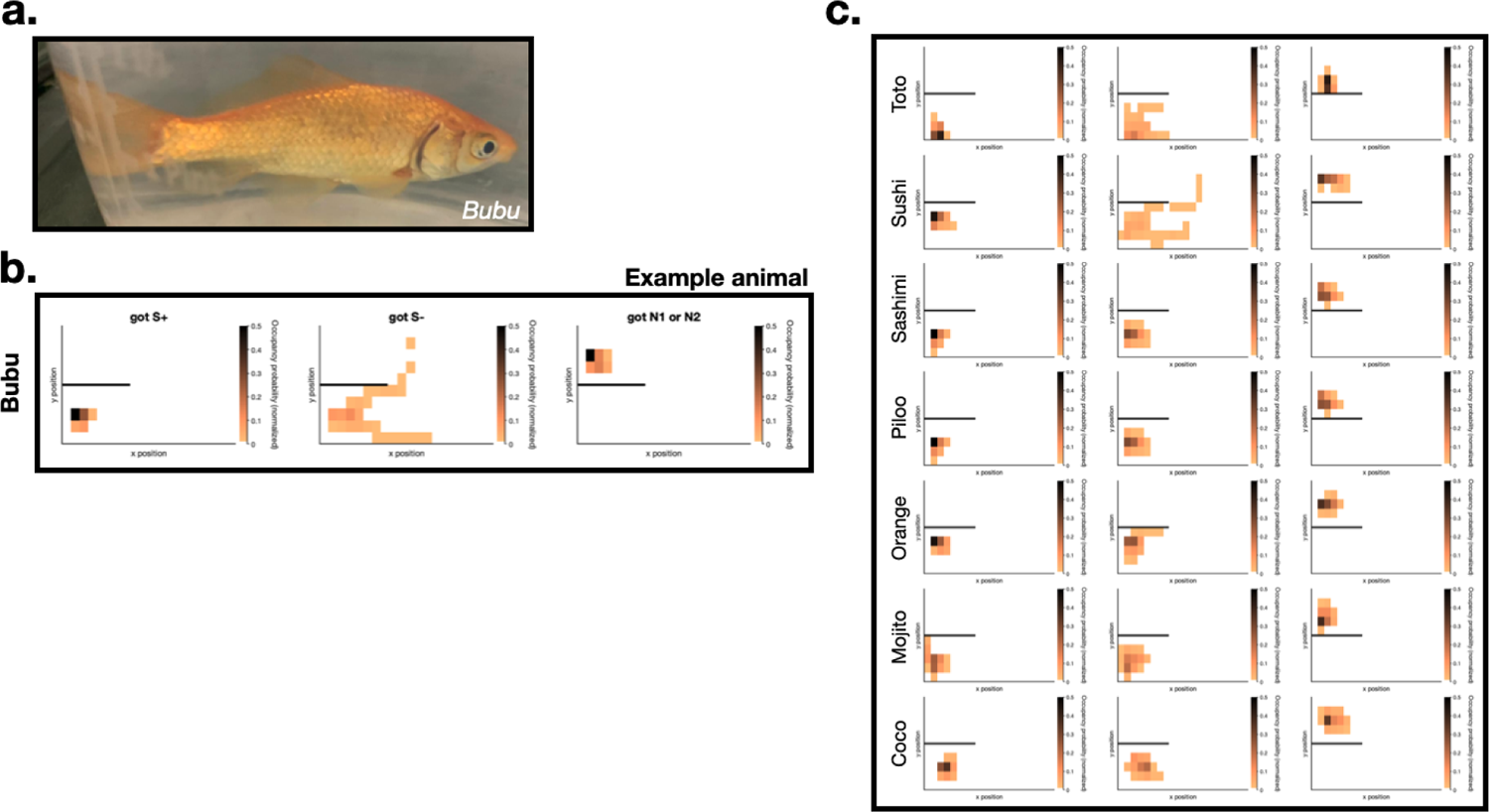
Subject’s post-choice occupancy reflects discrimination of stimuli signalling future outcomes. **a**. Example animal, *Bubu*, used for data in Figure 2. **b**. Normalised 2D histograms of occupancy averaged across 5 experimental days for the example animal (n=10 sessions) for stimuli S+, S-, and N1,N2, respectively. **c**. Same as in b. for the remaining 7 subjects (Sushi: 8; Toto: 7; Sashimi: 10; Piloo: 10; Orange: 10; Mojito: 8; Coco: 4 sessions, respectively).

**Supplementary Figure 3.**
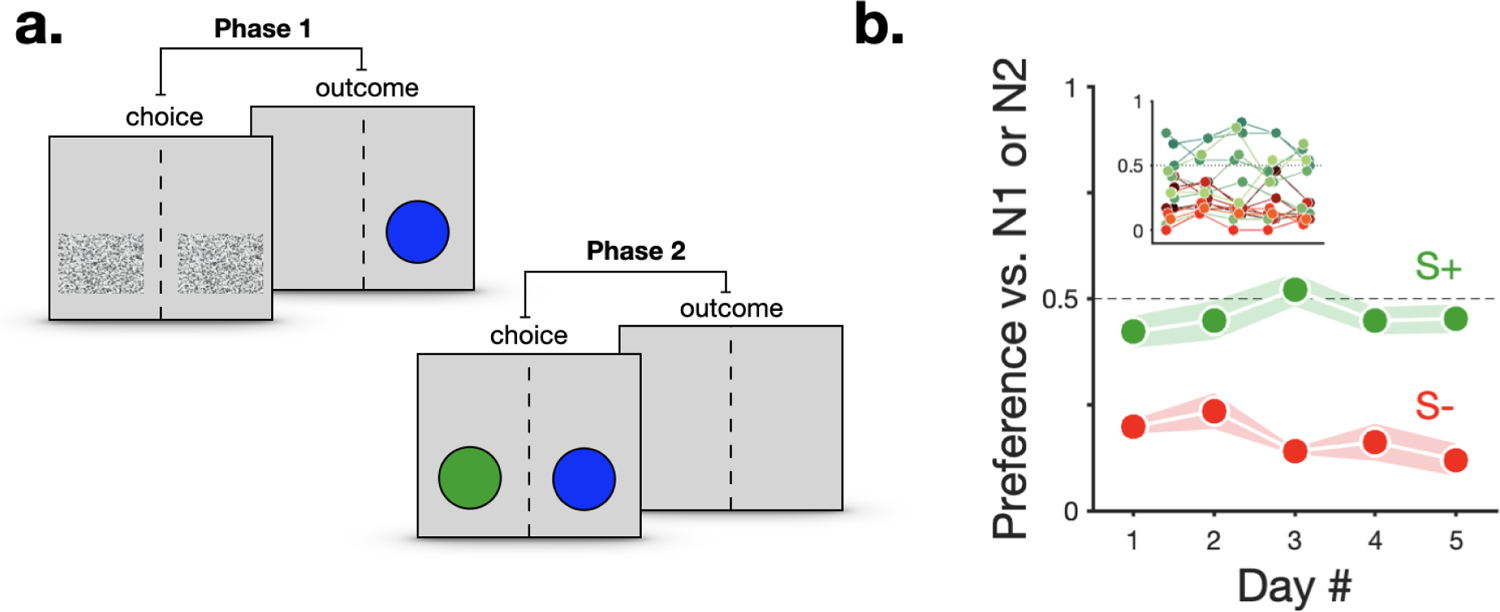
Goldfish preferences between reward-predictive stimuli. **a.** Rather than choosing between Info and NoInfo (*Phase 1*; **a**, left), in *Phase 2* (**a**, right) subjects were presented with the following pairs of outcome predictive stimuli to choose from: S+ (100% reward) vs. N1 or N2 (50% reward), and S- (0% reward) vs. N1 or N2 (50% reward). **b.** Proportion of choices for S+ (green) or S- (red) vs N1 or N2. Markers show group means and shading the s.e.m (n=8 animals). Inset shows individual means across 5 days.

---

**Supplementary Table 1.** Proportion of 1-option forced and 2-option choice trials across experimental days and experimental sessions from *Phase 1* of the experiment.

| Day | Session | Forced | Choices |
| --- | --- | --- | --- |
| 1-15 | 1 | 24 | -- |
|  | 2 | " | " |
| 16-20 | 1 | " | " |
|  | 2 | 16 | 8 |
| 21-40 | 1 | " | " |
|  | 2 | " | " |
| 41-45 | 1 | 8 | 16 |
|  | 2 | " | " |

**Supplementary Table 2.**
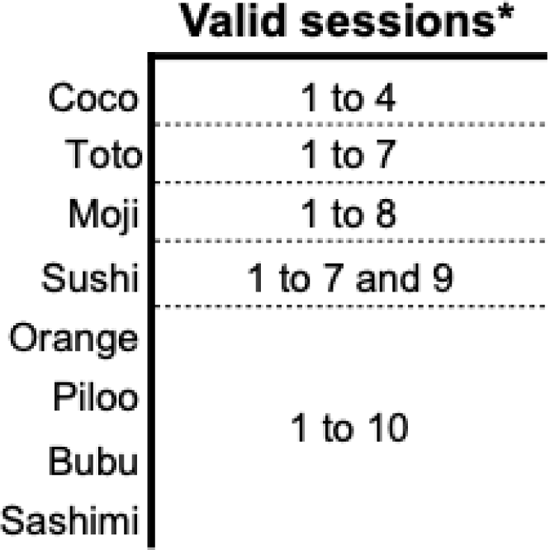
Number of valid sessions included in the analysis of subject’s movement during the presentation of outcome predictive stimuli in the last 5 days of *Phase 1* (*out of 10 possible sessions).

